# Proteomic analysis of peripheral nerve myelin during murine aging

**DOI:** 10.1101/2023.04.26.538413

**Authors:** Dario Lucas Helbing, Joanna M. Kirkpatrick, Michael Reuter, Julia Bischoff, Amy Stockdale, Annemarie Carlstedt, Emilio Cirri, Reinhard Bauer, Helen Morrison

**Affiliations:** Leibniz Institute on Aging, Fritz Lipmann Institute, 07745 Jena, Germany; Department of Psychiatry and Psychotherapy, Jena University Hospital, Friedrich Schiller University Jena, 07745 Jena; Center for Intervention and Research on adaptive and maladaptive brain Circuits underlying mental health (C-I-R-C), Jena-Magdeburg-Halle; Institute of Molecular Cell Biology, Jena University Hospital, Friedrich Schiller University Jena, 07745 Jena

**Author notes:** <u>Correspondence should go</u> to: Helen Morrison Leibniz Institute on Aging, Fritz Lipmann Institute Beutenbergstraße 11 07745 Jena, Germany. Email addresses of all authors: Dario Lucas Helbing Joanna M. Kirkpatrick Michael Reuter Julia Bischoff Amy Stockdale Annemarie Carlstedt Emilio Cirri Reinhard Bauer Helen Morrison.

**Keywords:** Peripheral nervous system, Aging, Myelin, Proteomics, Extracellular Matrix

## Abstract

Aging of the peripheral nervous system (PNS) is associated with structural and functional changes that lead to a reduction in regenerative capacity and the development of age-related peripheral neuropathy. Myelin is central to maintaining physiological peripheral nerve function and differences in myelin maintenance, degradation, formation and clearance have been suggested to contribute to age-related PNS changes. Recent proteomic studies have elucidated the complex composition of the total myelin proteome in health and its changes in neuropathy models. However, changes in the myelin proteome of peripheral nerves during aging have not been investigated. Here we show that the proteomes of myelin fractions isolated from young and old nerves show only subtle changes. In particular, we found that the three most abundant peripheral myelin proteins (MPZ, MBP and PRX) do not change in old myelin fractions. We also show a tendency for high-abundance myelin proteins other than these three to be downregulated, with only a small number of ribosome-related proteins significantly downregulated and extracellular matrix proteins such as collagens upregulated. In addition, we illustrate that the peripheral nerve myelin proteome reported in this study is suitable for assessing myelin degradation and renewal during peripheral nerve degeneration and regeneration. Our results suggest that the peripheral nerve myelin proteome is relatively stable and undergoes only subtle changes in composition during mouse aging. We proffer the resultant dataset as a resource and starting point for future studies aimed at investigating peripheral nerve myelin during aging. Said datasets are available in the PRIDE archive under the identifier PXD040719 (aging myelin proteome) and PXD041026 (sciatic nerve injury proteome).

## I. Introduction

Aging results in profound alterations in morphology, function and regenerative potential of the peripheral nervous system (PNS) (1). Particularly, nerve fibre loss and myelin abnormalities have been observed, accompanied by changes in Schwann cell activity and differentiation state (1, 2, 3). Advanced age heralds a marked decrease in the number of primarily myelinated fibres in mice (1), whilst myelin morphology itself is altered, indicating chronic degeneration: Several abnormalities have been shown to occur in aged mice, including folded myelin loops, separation of myelin lamellae, myelin degeneration and the presence of myelin debris in phagocytic Schwann cells and macrophages (1, 3, 4). At the same time, the breakdown and clearance of myelin by Schwann cells and macrophages is reduced in old mice following a peripheral nerve injury (PNI) (2, 4), suggesting a defective phagocytic activity, probably due to either cellular differences in old phagocytic cells or changes in the composition of old myelin – since myelin components themselves can regulate myelin phagocytosis (5). Yuan et al. demonstrated recently that age-related pathological changes in myelin may be related to dysfunctional macrophages, as depletion of macrophages in old mice led to an amelioration of myelin abnormalities and demyelination (4). To the best of our knowledge, it has not been investigated as to whether the composition of myelin itself changes during aging, thus contributing to age-related myelin pathology. In fact, in addition to insulating axons to allow saltatory conduction (6), myelin serves many other functions in the PNS and central nervous system (CNS): Myelin composition regulates macrophage efflux from peripheral nerves after an injury (7), axonal cytoskeleton and glia-to-axon molecule transfer (8). Differences in the lipid or protein composition of PNS and CNS myelin due to genetic mutations have been linked to a variety of human neurological diseases (6, 8). Multiple studies have also investigated myelin turnover and differences in the myelin composition in primarily CNS pathologies. On the one hand, along with nuclear pore proteins, myelin proteins are thought to be among the longest-lived proteins in the murine brain (9, 10). Accordingly, it has been shown that CNS myelin in humans is subject to slow but lifelong turnover (11). On the other hand, evidence from several human diseases and corresponding animal models indicates changes in myelin composition, suggesting differences in myelin synthesis and/or maintenance: For example, peripheral myelin proteome analysis in the Prnp0/0 mouse model of chronic demyelinating polyneuropathy identified SEPT9 as a potential myelin-cytoskeleton associated protein that was upregulated compared to wildtype myelin (12). Recently, the same research group presented the most accurate quantification to date of the peripheral mouse myelin proteome (13). This study confirmed their earlier findings that myelin protein zero (P0/MPZ), myelin basic protein (MBP), and periaxin (PRX) account for more than 75% of the entire murine PNS myelin (13). Moreover, the authors obtained proteome profiles from Prx-/- mice, another mouse model of peripheral neuropathy, specifically Charcot-Marie-Tooth disease type 4F. It was observed that the myelin proteome showed significant remodeling, in particular extracellular matrix (ECM) proteins and immune system-related proteins were differentially abundant in the myelin fraction isolated from neuropathic mice (13). Taken together, morphological changes in myelin occur during both aging and peripheral neuropathy, with prior research showing remodeling of the myelin proteome in the latter. Previous studies have also demonstrated a decrease of major myelin proteins in aged mice in total nerve lysates; this is mitigated by caloric restriction, a known neuroprotective intervention (14, 15). However, isolated PNS myelin has not been studied during aging to determine differences in myelin composition. Thus, the focus of this study was to conduct an unbiased comparison of proteins associated with isolated PNS myelin in young (2-3 months) and old mice (18 months). Moreover, we correlate our quantitative proteome data with the PNS myelin proteome data determined by Siems et al. (13) to identify commonly found “PNS core myelin proteins”. The validity of the myelin proteome profiles obtained is exemplified by the observation that the majority of detected proteins are profoundly downregulated during PNS degeneration in whole sciatic nerve proteomes acquired at different time points after a nerve crush injury. Functional enrichment analyses reveal primarily ribosome and ECM changes within aging PNS myelin. Potential implications of myelin-associated ribosomal and ECM-/collagen changes for peripheral nervous system aging will be further discussed.

## II. Materials and Methods

### Experimental animals

All animals were on a C57BL/6 J background. Mice were group housed, with free access to standard chow and water and maintained with 12 h light and dark cycles. Temperature was 20 ± 2 °C during the experimental period. The animal procedures were performed according to the guidelines from Directive 2010/63/EU of the European Parliament on the protection of animals used for scientific purposes. Experiments were approved by the Thuringian State Office for Food Safety and Consumer Protection (license FLI-17-007). Mice were considered to be old at around 18 months and considered young at around 2-3 months.

### Sciatic nerve crush injury

Unilateral injuries of sciatic nerves were performed with minimal invasion, as described previously (16, 17). Briefly, mice were anaesthetized with isoflurane in oxygen, fur was removed with an electric razor (Aesculap ISIS, B. Braun AG, Melsungen Germany) and skin incised minimally. The biceps femoris muscle was separated to reveal the sciatic nerve. Using a smooth hemostatic forceps (width 0,6 mm, curved, clamping length 19 mm, Fine Science Tools GmbH, Heidelberg, Germany) the nerve was crushed mid-thigh. Muscle tissue was sutured using non-absorbable surgical suture material (Catgut GmbH, Markneukirchen, Germany) and skin closed with an AutoClip System (FST, 9 mm clips).

### Immunohistochemistry

Sciatic nerves were fixed in 4% PFA for 24 h at 4 °C and incubated for at least 24 h in PBS at 4 °C. Afterwards, the nerves were transferred into biopsy embedding cassettes and dehydrated in 30% EtOH for at least 30 min before paraffin embedding. 5 μm paraffin sections were cut at a microtome Microm HM355S using Microm SEC 35 blades. Sections were deparaffinized and rehydrated and epitope retrieval performed using 10 mM sodium citrate buffer, pH6 at 80 °C for 30 min. Blocking was performed at room temperature (RT) for 1 h 45 min in blocking buffer (0.2% fish skin gelatin, 2% donkey serum, 1% BSA and 0.4% Triton-X in PBS). Primary antibodies were diluted 1:50 (Col4a1, NBP1-26549) and 1:100 (MBP, NB-600-717) in blocking buffer and incubated overnight at 4 °C. After multiple washing steps, sections were incubated with secondary antibodies (donkey anti-rat Alexa488 (Thermo Fisher Scientific, A21208) and donkey anti-rabbit Alexa647 (Thermo Fisher Scientific, A31573)) at room temperature for 2 h. Sections were washed, DAPI-stained and mounted with glass coverslips in ImmuMount™ (Thermo Fisher Scientific) and imaged on an Axio Scan.Z1 device with with a 20x objective and ZEN software. All fluorescence intensity quantifications were semi-automatic with Fiji (ImageJ version 1.53t)(18) using custom macros. Quantification of Col4a1 fluorescence intensity was achieved by measuring the raw integrated density within Col4a1^+^ area and normalization to the Col4a1^+^ area. Quantification of Col4a1 myelin coverage was done through extraction of MBP^+^ area into a mask and measurement of the Col4a1^+^ area within this MBP mask. Lastly, Col4a1 fluorescence intensity in overlap to myelin was measured through quantification of the raw integrated density of Col4a1 fluorescent signal and subsequent normalization to the percentage of the Col4a1^+^ area within MBP mask, as well as the MBP^+^ area.

### Myelin Isolation

Myelin was isolated from pooled sciatic nerve fractions (n = 7-10 nerves per pool) with one pool representing one biological replicate (n = 3 biological replicates for each age), as described previously (19). In brief, sciatic nerves were isolated from mice on a C57BL/6 J background and snap-frozen in liquid nitrogen. Pooled sciatic nerves were lysed in 0.27 M Sucrose using a Polytron homogenizer and carefully added to ultracentrifuge tubes containing 0.83 M sucrose. Subsequently, samples were ultra-centrifuged for 45 min at 82,000 x g. The formed myelin intermediate phase was isolated, washed in TRIS-Cl buffer, centrifuged again for 15 min at 82,000 x g and resuspended in TRIS-Cl buffer. Then, myelin protein pellets were centrifuged again for 10 min at 17,000 x g, before samples were resuspended in PBS and frozen at -80 °C until further proteomic characterisation.

### Sample preparation and LC-MS/MS for myelin proteome samples

The myelin pellets were resuspended in lysis buffer (final concentration: 0.1 M HEPES/pH 8; 2 % SDS; 0.1 M DTT), vortexed, then sonicated using a Bioruptor (Diagenode) (10 cycles of 1min on and 30s off with high intensity @ 20°C). For reduction and full denaturation of the proteins, the lysates were first incubated at 95 °C for 10 min and subsequently sonicated in the Bioruptor for a further 10 cycles, as before. The supernatants (achieved after centrifugation, 14,000 rpm, room temperature, 2 minutes) were then treated with iodacetamide (room temperature, in the dark, 30 min, 15 mM). After running a coomassie gel to estimate the amount of protein in each sample, approximately 60-120 µg of each sample were treated with 4 volumes ice-cold acetone to 1 volume sample and left overnight at -20 °C, to precipitate the proteins. The samples were then centrifuged at 14,000 rpm for 30 min, 4 °C. After removal of the supernatant, the precipitates were washed twice with 400 µL 80% acetone (ice-cold). Following each wash step, the samples were vortexed, then centrifuged again for 2 minutes at 4 °C. The pellets were allowed to air-dry before dissolution in digestion buffer (60 µL, 3 M urea in 0,1 M HEPES, pH 8) with sonication (3 cycles in the Bioruptor as above) and incubation for 4 h with LysC (1:100 enzyme: protein ratio) at 37 °C, with shaking at 600 rpm. The samples were then diluted 1:1 with milliQ water (to reach 1.5 M urea) and incubated with trypsin (1:100 enzyme: protein ratio) for 16 h at 37 °C. The digests were acidified with 10% trifluoroacetic acid, then desalted with Waters Oasis® HLB µElution Plate 30 µm in the presence of a slow vacuum. This process involved conditioning the columns with 3×100 µL solvent B (80% acetonitrile; 0.05% formic acid) and equilibration with 3×100 µL solvent A (0.05% formic acid in milliQ water). The samples were loaded, washed 3 times with 100 µL solvent A, then eluted into PCR tubes with 50 µL solvent B. The eluates were dried down with the speed vacuum centrifuge and dissolved in 50 µL 5% acetonitrile, 95% milliQ water, with 0.1% formic acid prior to analysis by LC-MS/MS.

For LC-MS/MS, peptides were separated using the nanoAcquity UPLC system (Waters) fitted with a trapping (nanoAcquity Symmetry C18, 5 µm, 180 µm x 20 mm) and an analytical column (nanoAcquity BEH C18, 1.7 µm, 75 µm x 250 mm). The outlet of the analytical column was coupled directly to an Orbitrap Fusion Lumos (Thermo Fisher Scientific) using the Proxeon nanospray source. Solvent A was water, 0.1 % formic acid and solvent B was acetonitrile, 0.1 % formic acid. The samples (500 ng) were loaded onto the trapping column for 6 minutes, with a constant flow of solvent A at 5 µL/min. Peptides were eluted via the analytical column with a constant flow of 0.3 µL/min. During this step, the percentage of solvent B increased in a linear fashion from 3 % to 25 % in 30 minutes, then to 32 % in 5 more minutes and finally to 50 % in a further 0.1 minutes. Total runtime was 60 minutes. The peptides were introduced into the mass spectrometer via a Pico-Tip Emitter 360 µm OD x 20 µm ID; 10 µm tip (New Objective) and a spray voltage of 2.2 kV applied. The capillary temperature was set at 300 °C. and the RF lens to 30%. Full scan MS spectra with mass range 375-1500 m/z were acquired in profile mode in the Orbitrap with a resolution of 120,000. The filling time was set at maximum of 50 ms with limitation of 2 x 10^5^ ions. The “Top Speed” method was employed to take the maximum number of precursor ions (with an intensity threshold of 5 x 10^3^) from the full scan MS for fragmentation (using HCD collision energy, 30%) and quadrupole isolation (1.4 Da window) and measurement in the ion trap, with a cycle time of 3 seconds. The MIPS (monoisotopic precursor selection) peptide algorithm was employed, but with relaxed restrictions when too few precursors meeting the criteria were found. The fragmentation was conducted after accumulation of 2 x 10^3^ ions, or after filling time of 300 ms for each precursor ion (whichever occurred first). MS/MS data were acquired in centroid mode, with the Rapid scan rate and a fixed first mass of 120 m/z. Only multiply charged (2+ - 7+) precursor ions were selected for MS/MS. Dynamic exclusion was employed with maximum retention period of 60 seconds and relative mass window of 10 ppm. Isotopes were excluded. Only 1 data dependent scan was performed per precursor (only the most intense charge state selected). Ions were injected for all available parallelizable time. A lock mass correction using a background ion (m/z 445.12003) was applied to improve mass accuracy. Data acquisition was performed using Xcalibur 4.0/Tune 2.1 (Thermo Fisher Scientific).

### Mass spectrometry data analysis for myelin proteome samples

For the quantitative label-free analysis, raw files from the Orbitrap Fusion Lumos were analyzed using MaxQuant (version 1.5.3.28) (20). MS/MS spectra were searched against the Mus musculus Swiss-Prot entries of the Uniprot KB (database release 2016_01, 16,755 entries) using the Andromeda search engine (21). A list of common contaminants was appended to the database search with search criteria set as follows: full tryptic specificity was required (cleavage after lysine or arginine residues, unless followed by proline); 2 missed cleavages were allowed; oxidation (M) and acetylation (protein N-term) were applied as variable modifications, with mass tolerances of 20 ppm set for precursor and 0.5 Da for fragments. The reversed sequences of the target database were used as decoy database. Peptide and protein hits were filtered at a false discovery rate of 1% using a target-decoy strategy (22). Additionally, only proteins identified by at least 2 unique peptides were retained. The intensity Based Absolute Quantification values (iBAQ, which corresponds to the sum of all the peptides intensities divided by the number of observable peptides of a protein, from the proteinGroups.txt output of MaxQuant) were used for further analysis. All comparative analyses were performed using R version 3.2.3. (23) – the R packages MSnbase (24) for processing proteomics data and the included package imputeLCMD for imputing missing values based on the definitions for values MAR and MNAR. The latter were defined for each pairwise comparison as values that were (i) missing in 4 or 5 out of 5, or 3 out of 5 biological replicates in one sample group, and (ii) present in 3 of the 5 biological replicates in the second sample group. Given their non-random distribution across samples, these values were considered as underlying biological differences between sample groups. MNAR values were computed using the “MinDet” method, by replacing values with minimal values observed in the sample. MAR were consequently defined for each pairwise comparison as values that were missing in 2 out of 5 biological replicates per sample group. MAR values were imputed based on the “knn” method (k-nearest neighbors) (24). All other cases (e.g., protein groups with less than 2 values in both sample groups) were filtered out due to lack of sufficient information to perform robust statistical analysis. The data was quantile normalized to reduce technical variations. Protein differential expression was evaluated using the Limma package (25). Differences in protein abundances were statistically determined using the Student’s t-test with variances moderated by Limma’s empirical Bayes method. False discovery rate was estimated using fdrtool (26). The obtained dataset was used for further analyses with R (23) and has been deposited in the PRIDE repository (PXD040719) (27).

### RNA and protein isolation from sciatic nerve tissue

Frozen sciatic nerves, 1.4 mm ceramic (zirconium oxide) beads and peqGOLD TriFast™ were placed in a 2 ml tube and homogenized using a PeqLab-Precellys® 24 homogenizer. Samples were transferred to RNA-free tubes, mixed with 0.25 ml chloroform for 10-15 sec and further incubated for 5 min at RT. Centrifugation for 15 min at 12,000 x g at 4 °C produced a separation of two phases; the upper phase containing RNA and the lower phase containing the protein fraction. Proteins were obtained from the protein fraction by adding two volumes of isopropanol. Mixed solution was incubated for 10 min at RT or on ice, followed by 10 min centrifugation at 12,000 x g at 4 °C. The pellet was washed twice with 95 % ethanol before transfer to the FLI Core Facility Proteomics for further experiments.

### Proteomics of injured sciatic nerves

#### Sample preparation

For proteomics analysis, crushed nerve samples were resuspended in 100 µl PBS and lysis buffer (fc 4% SDS, 100 mM HEPES, pH 8.5, 50 mM DTT), then sonicated in a Bioruptor (Diagenode, Beligum) (10 cycles of 1 minute on and 30s off with high intensity @ 20°C). Next, samples were heated at 95 °C for 10 min, before another round of sonication in the Bioruptor. The lysates were clarified and debris precipitated by centrifugation at 14,000 rpm for 10 min, then incubated with iodacetamide (room temperature, in the dark, 20 min, 15 mM). 10 µg of the sample was removed to check lysis on a coomassie gel. Based on the gel, an estimated 50 µg of each sample was treated with 8 volumes of ice-cold acetone and left overnight at -20 °C to precipitate the proteins. These samples were then centrifuged at 14,000 rpm for 30 minutes, 4 °C. Following removal of the supernatant, the precipitates were washed twice with 300 µL of a solution of ice-cold 80 % acetone. After addition of each wash solution, the samples were vortexed and centrifuged again for 10 min at 4°C. The pellets were allowed to air-dry before dissolution in digestion buffer at 1 µg/µL (1 M guanidine HCl in 0.1 M HEPES, pH 8). To facilitate resuspension of the protein pellet, the samples were subjected to 5 cycles of sonication in the Bioruptor, as described above. Afterwards, LysC (Wako) was added at 1:100 (w/w) enzyme:protein ratio and digestion proceeded for 4 h at 37 °C under shaking (1000 rpm for 1 h, then 650 rpm). The samples were diluted 1:1 with milliQ water (to reach 1.5 M urea) and incubated with a 1:100 w/w amount of trypsin (Promega sequencing grade) overnight at 37 °C, 650 rpm. Next, the digests were acidified with 10% trifluoroacetic acid before desalting with Waters Oasis® HLB µElution Plate 30 µm in the presence of a slow vacuum. In this process, the columns were conditioned with 3×100 µL solvent B (80% acetonitrile; 0.05% formic acid) and equilibrated with 3x 100 µL solvent A (0.05% formic acid in milliQ water). The samples were loaded, washed 3 times with 100 µL solvent A, then eluted into PCR tubes with 50 µL solvent B. The eluates were dried down with the speed vacuum centrifuge and dissolved in 5% acetonitrile, 95% milliQ water, with 0.1% formic acid at a concentration of 1 µg/µL. 10 µL was transferred to an MS vial and 0.25 µL of HRM kit peptides (Biognosys, Zurich, Switzerland) was spiked into each sample prior to analysis by LC-MS/MS.

#### LC-MS/MS data indipendent acquisition (DIA)

Peptides were separated using the M-Class nanoAcquity UPLC system (Waters) fitted with a trapping (nanoAcquity Symmetry C18, 5 µm, 180 µm x 20 mm) and an analytical column (nanoAcquity BEH C18, 1.7 µm, 75 µm x 250 mm). The outlet of the analytical column was coupled directly to a Thermo Q-Exactive HFX using the Proxeon nanospray source. Solvent A was water, 0.1 % formic acid and solvent B was acetonitrile, 0.1 % formic acid. The samples (approx. 1 µg) were loaded with a constant flow of solvent A, at 5 µL/min onto the trapping column for 6 minutes. Peptides were eluted via a non-linear gradient from 1% to 62.5% B in 131 min. Total runtime was 145 min, including clean-up and column re-equilibration. The peptides were introduced into the mass spectrometer via a Pico-Tip Emitter 360 µm OD x 20 µm ID; 10 µm tip (New Objective) and a spray voltage of 2.2 kV applied. The RF ion funnel was set to 40%.

The conditions for DDA were as follows: Full scan MS spectra with mass range 350-1650m/z were acquired in the Orbitrap with a resolution of 60,000 FWHM. The filling time was set at maximum of 20ms with an AGC target of 3×10^6^ ions. A Top15 method was employed to select precursor ions from the full scan MS for fragmentation, quadrupole isolation (1.6m/z) and measurement in the Orbitrap (resolution 15,000 FWHM, fixed first mass 120m/z). The fragmentation was conducted after accumulation of 2×10^5^ ions, or after filling time of 25 ms for each precursor ion (whichever occurred first). Only multiply charged (2+ -7+) precursor ions were selected for MS/MS. Dynamic exclusion was employed with maximum retention period of 20 s. Isotopes were excluded.

For DIA analysis MS conditions were set up as follows: Full scan MS spectra with mass range 350-1650 m/z were acquired in profile mode in the Orbitrap, with a resolution of 120,000 FHWM. The filling time was set at maximum of 60 ms with limitation of 3 x 10^6^ ions. DIA scans were acquired with 40 mass window segments of differing widths across the MS1 mass range. HCD fragmentation (stepped normalized collision energy; 25.5, 27, 30%) was applied and MS/MS spectra were acquired with a resolution of 30,000 FHWM, with a fixed first mass of 200 m/z after accumulation of 3 x 10^6^ ions or after filling time of 40 ms (whichever occurred first). Data were acquired in profile mode. Tune version 2.9 and Xcalibur 4.1 were employed for data acquisition and processing.

#### Data analysis

Acquired data were processed using Spectronaut Professional v17 (Biognosys AG). For library creation, the DDA and DIA raw files were searched with Pulsar (Biognosys AG) against the human Swiss-Prot database (Mus musculus, entry only, release 2016_01, 16,747 entries) with a list of common contaminants appended, using default settings. Default BGS factory settings were used for library generation. Data were searched with the following modifications: Carbamidomethyl (C) (fixed) and Oxidation (M), Acetyl (Protein N-term). A maximum of 2 missed cleavages for trypsin and 5 variable modifications were allowed. Identifications were filtered to achieve an FDR of 1 % at the peptide and protein levels.

For DIA analysis, relative quantification was performed in Spectronaut for each pairwise comparison, using replicate samples from each condition with default settings, except: Major Group Quantity = median peptide quantity; Major Group Top N = OFF; Minor Group Quantity = median precursor quantity; Minor Group Top N = OFF; Data Filtering = Q value; Normalization Strategy = Local normalization; Row Selection = Automatic; Exclude Single Hit Proteins = TRUE. Tests for differential abundance were performed using a paired t-test between intact and crushed nerve from each mouse. P values were corrected for multiple testing using the method described by Storey (28) to obtain false discovery rate adjusted p values (q values). The obtained dataset was used for further analyses with R (23) and has been deposited in the PRIDE repository (PXD041026) (27).

#### Proteomic and bioinformatic data analysis

All data analysis steps were completed with R (23). Data wrangling steps were performed using primarily dplyr functions from the tidyverse package (29). Only proteins detected in at least 2 out of 3 myelin pools within each biological group (young or old) were considered in all further analyses. Venn diagrams and heatmaps aside, all plots were created with custom code using the ggplot2 package within the tidyverse package (29). Correlation analyses were achieved using the cor.test function from the R stats package and depicted correlation coefficients represent Spearman’s ρ as in all cases data were not normally distributed. Gene ontology annotations for cellular compartments were retrieved using the biomaRt package (30) as previously described (31). Venn diagrams were made using the VennDiagram package (32). Comparative analysis to the myelin proteome determined by Siems et al. (13) was performed using the PPM values from the MS^e^ proteome (Source data 1 associated with Figure 1 from Siems et al.). For heatmap generation of proteomics data from nerves post-injury, protein group quantities as reported by Spectronaut 17 were log_2_ transformed and subsequently row-wise z-scored across all samples. Heatmaps were created using the ComplexHeatmap package (33) and reordering of the dendrogram using the dendsort package (34). Reactome pathway enrichment analyses were performed using DAVID (35) with the whole myelin proteome determined in this study as a background.

**Figure 1.**
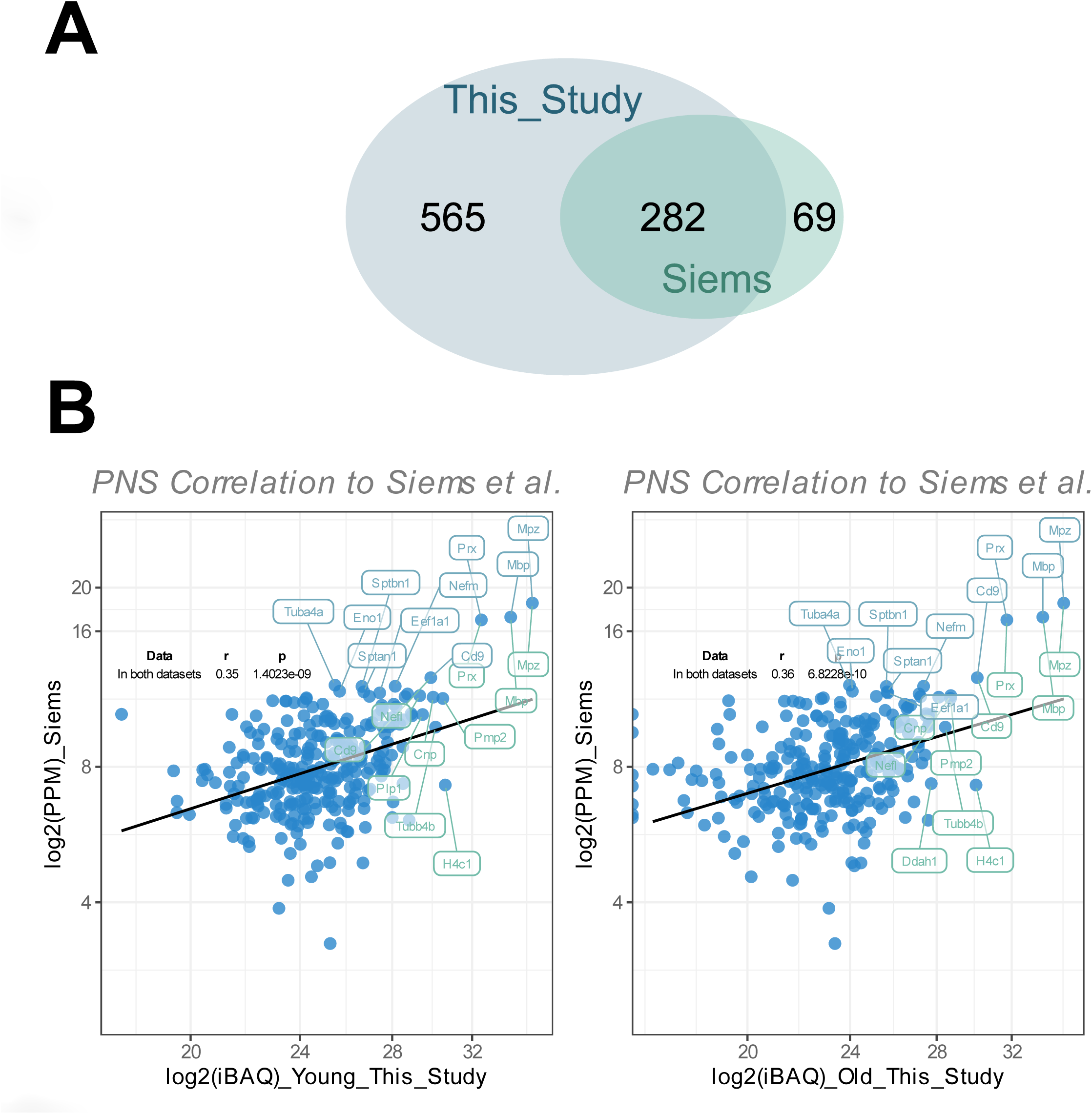
**A** Venn diagram illustrating the number of myelin proteins detected in this study compared to that of Siems et al. (13). **B** Correlation plot of the mean Log_2_(iBAQ-values) of detected myelin proteins in young (left figure) or old nerves (right figure) to the mean PPM (= parts per million) values of the MS^e^-obtained myelin proteome dataset of young, wildtype mice from the Siems et al. study. The top 10 proteins with the highest mean iBAQ-values in each young or old myelin fraction, as well as the top 10 proteins for the Siems et al. myelin proteome data, are labelled with their corresponding gene names (top 10 young/old proteins in green, top 10 Siems et al. proteins in turquoise). Correlation statistics are shown in the upper left corner.

#### Statistical procedures

All analyses were performed using Graphpad Prism 8.4 or R (23). Shapiro-Wilk normality test was carried out before further statistical analysis was performed. Depending on the result, parametric (Unpaired, two-tailed Student’s t-test) or non-parametric (Friedman-test with post-hoc Dunn’s multiple comparison test) statistical tests were conducted, as stated in the figure legends. Statistical tests for proteomic analysis and functional annotation and enrichment analyses were performed as indicated in the respective material and methods section. Effect size (Cohen’s d) was calculated using G*Power v3.1.9.1 (36). All data are presented as mean ± SEM. Statistical significance was accepted if the p-value (p) was p ≤ 0.05 (depicted as “*” or “#”).

## III. Results

### Comparison to a previously reported PNS myelin proteome

Before performing any analysis focusing on proteome myelin changes between young and old nerves, we wanted to assess the validity of our dataset. As such, we compared the protein quantities and general overlap of the PNS myelin proteome obtained in this study with the currently most accurate PNS myelin proteome dataset of sciatic nerve myelin isolated from young, C57Bl/6 mice, as determined by Siems et al. (13). The majority of the proteins identified by Siems et al. were also identified in our proteomics experiment (282 out of 351 proteins, **Figure 1A**). However, we detected a significant number of potential PNS myelin proteins that were not detected in the MS^e^ proteome by the Siems group (565 out of a total of 847 proteins detected, **Figure 1A**). In addition, we plotted the protein quantities of all overlapping proteins and detected a significant positive correlation between protein quantities from both datasets (Spearman’s ρ = 0.35 and 0.36 for young and old myelin fractions, respectively, **Figure 1B**). Within the top 10 most abundant myelin proteins, 4 proteins were shared (MPZ, MBP, Prx and CD9), while other classical myelin proteins were detected, such as myelin protein P2 (PMP2), CNPase (CNP) and myelin proteolipid protein (PLP1) (**Figure 1B**).

### Proteomic downregulation of myelin proteins after sciatic nerve crush injury

Myelin is degraded after a peripheral nerve injury mainly by Schwann cells and macrophages (37, 38, 39, 40). As efficient myelin clearance is critical for nerve de- and subsequent nerve regeneration (2, 41, 42), we reasoned that we could further assess the validity of our dataset by examining the regulation of myelin proteins after a sciatic nerve injury. To this end, we performed proteomics from whole nerves, both from crushed and contralateral intact nerves 7 days and 28 days post crush injury (dpc), for comparative analyses to the obtained myelin proteome dataset. As seen in **Figure 2A**, we found that the majority of myelin proteins (∼71% at 7 dpc) were significantly downregulated after PNI (q < 0.05), which is greater than anticipated (ground probability for a protein to be downregulated (Log_2_FC < 0 and q < 0.05) 7 days crush vs. intact: 0.349 and 28 days crush versus intact: 0.408, binomial test 7 days crush vs. intact myelin proteins p = 4.346e-80, binomial test 28 days crush versus intact myelin proteins p = 1.043e-38). Typically, at 28 days after injury nerve regeneration and remyelination of regenerating axons is quite advanced (17). We also observed that myelin proteins that were downregulated at 7dpc, were less downregulated at 28dpc (**Figure 2A** and **B**). The protein abundance of degraded myelin proteins was significantly higher in crushed nerves 28dpc, as compared to the 7dpc cohort (**Figure 2B**), reflecting nerve regeneration and remyelination. Likewise, we tested whether overlapping myelin proteins detected in this study and by Siems et al. demonstrate the same pattern. Indeed, we observed that the majority of overlapping myelin proteins (∼76%) were also strongly downregulated at 7dpc (Log_2_FC < 0 and q < 0.05), which was also more than expected (same ground probabilities as in Figure 2, binomial test 7dpc crush versus intact overlapping myelin proteins p = 6.185e-37, binomial test 28dpc crush versus intact overlapping myelin proteins p = 1.546e-25) (**Figure S1A, B**). The protein abundancies of this subset were also elevated in crushed nerves 28dpc as compared to 7dpc crushed nerves (**Figure S1B**). The similar patterns of both datasets imply that the majority of myelin proteins reported in this study are most likely actually true myelin proteins.

**Figure 2.**
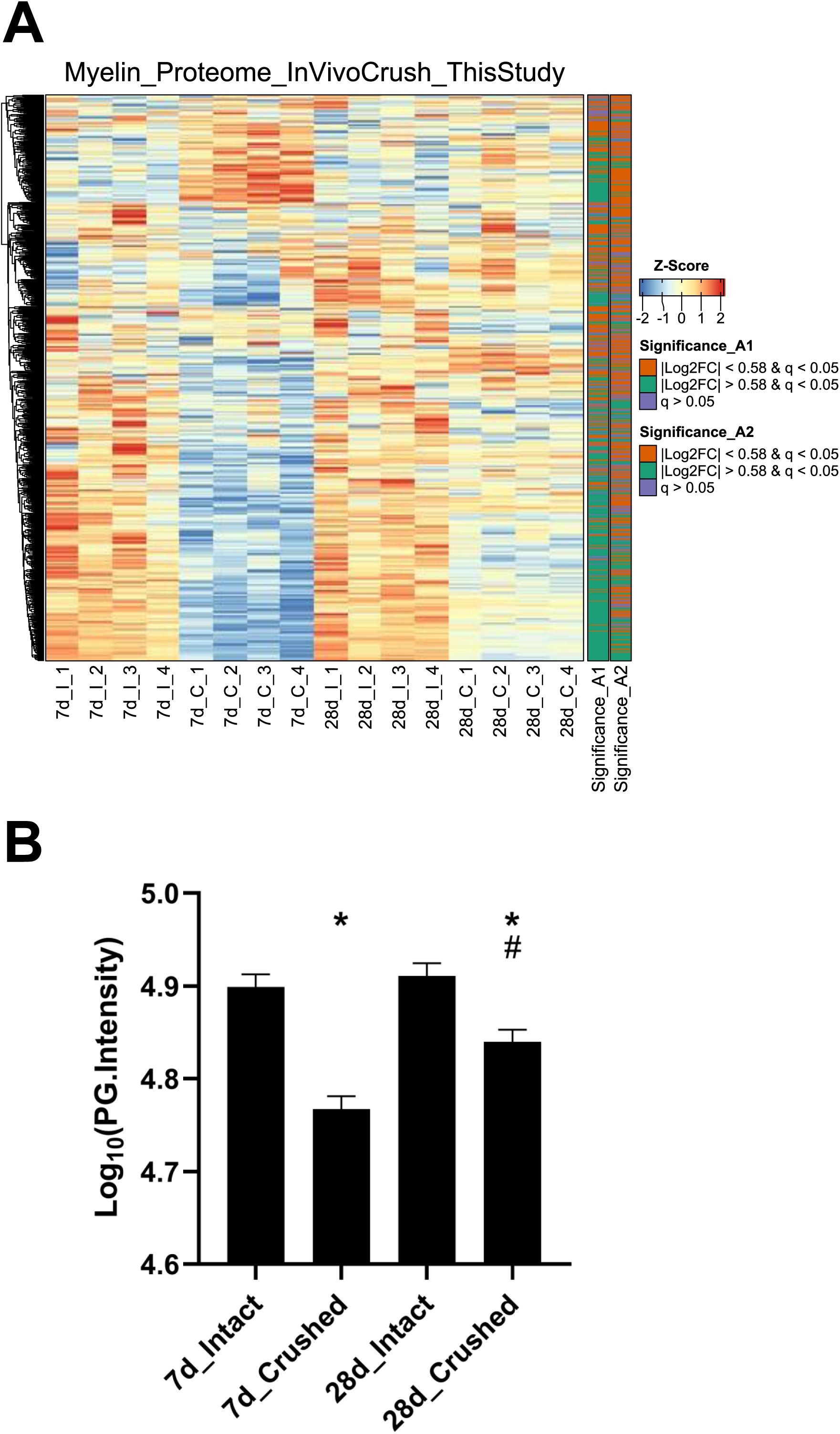
**A** The myelin proteome obtained in this study was mapped to a proteomics dataset of peripheral nerve degeneration and regeneration, in which sciatic nerve proteomes were obtained at different timepoints post crush injury. “7d_I” refers to contralateral, intact control nerves from the same mice as the “7d_C” groups that refer to the crushed, ipsilateral nerves 7 dpc, n = 4 mice per group. Similarly, the “28d_I” and “28d_C” groups represent the intact, contralateral and crushed, ipsilateral nerves from the same mice in each case 28 dpc, n = 4 mice per group. Only the proteins are shown that were changed with a q-value < 0.05 in either the comparison 7d crush versus 7d intact (abbreviated as “A1”), or the comparison 28d crush versus 28d intact (abbreviated as “A2”). The protein-wise z-score normalized protein group intensities of all mapped myelin proteins are shown; on the right side of the heatmap an annotation of Log_2_ FC and q-value levels are depicted (│Log_2_ FC│≥ 0.58 in green or │Log_2_ FC│≤ 0.58 in orange, meaning more or less than 50% change and q-value < 0.05, respectively or a q-value > 0.05 in purple). **B** Bar charts of Log_10_(PG.Intensities) of all 7dpc or 28dpc differentially expressed myelin proteins are shown (n = 699 proteins). A Friedman-test was conducted, with a p-value ≤ 0.05. Post-hoc Dunn’s multiple comparison tests were conducted with * = p ≤ 0.05 for comparisons 7d crush versus intact and 28d crush versus intact and # = p ≤ 0.05 for the comparison 28d crush versus 7d crush.

**Figure S1.**
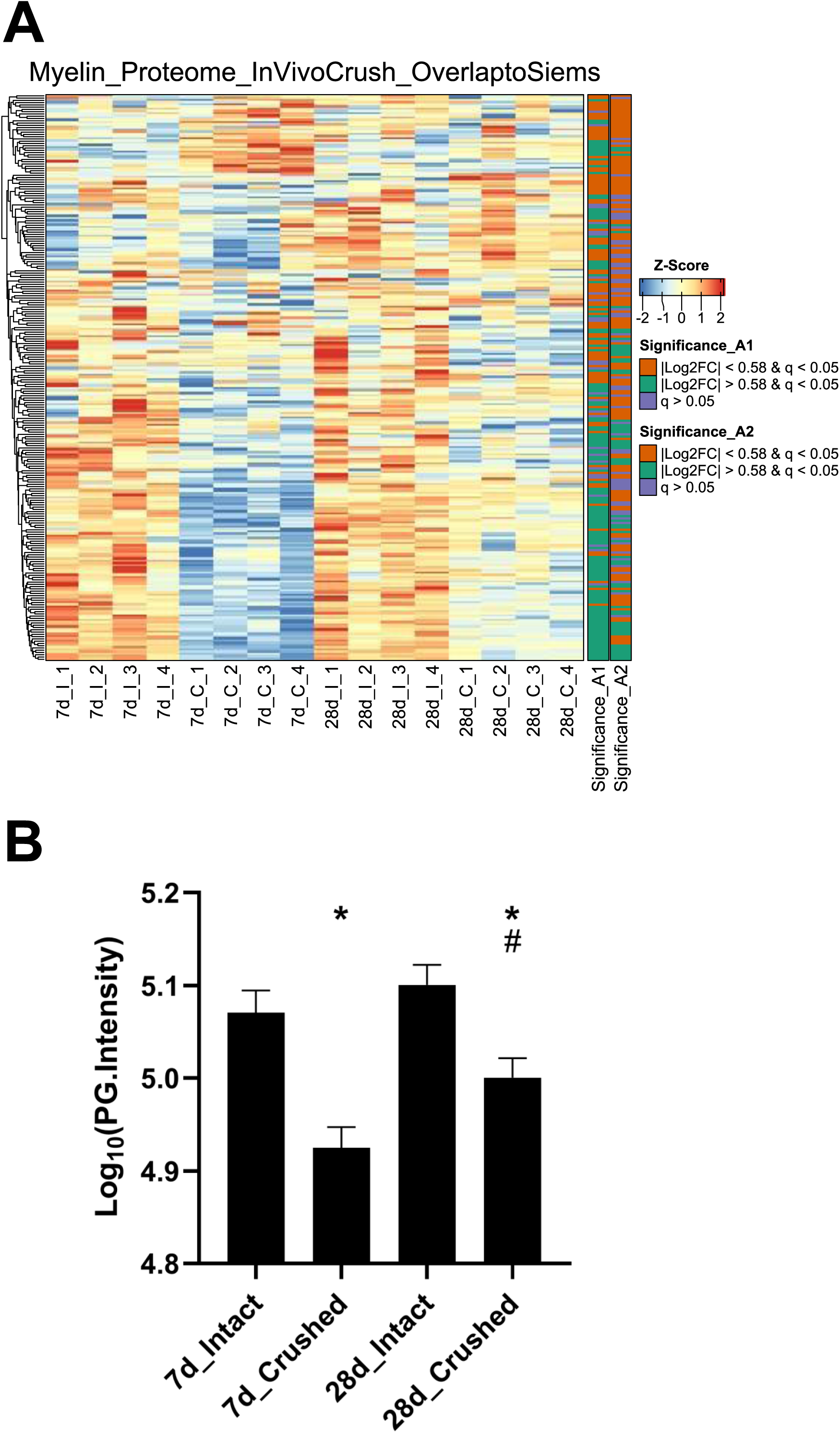
**A** The myelin proteome determined by Siems et al. was mapped to the proteomics dataset of peripheral nerve degeneration and regeneration as shown in figure 2. Equivalently, “7d_I” refers to contralateral, intact control nerves from the same mice as the “7d_C” groups that refer to the crushed, ipsilateral nerves 7 dpc, n = 4 mice per group. The “28d_I” and “28d_C” groups represent the intact, contralateral and crushed, ipsilateral nerves from the same mice in each case 28 dpc, n = 4 mice per group. Only the proteins are shown that were changed with a q-value < 0.05 in either the comparison 7d crush versus 7d intact (abbreviated as “A1”), or the comparison 28d crush versus 28d intact (abbreviated as “A2”). The protein-wise z-score normalized protein group intensities of all mapped myelin proteins are shown; on the right side of the heatmap an annotation of Log_2_ FC and q-value levels are depicted (│Log_2_ FC│≥ 0.58 in green or │Log_2_ FC│≤ 0.58 in orange, meaning more or less than 50% change and q-value < 0.05, respectively or a q-value > 0.05 in purple). **B** Bar charts of Log_10_(PG.Intensities) of all 7dpc or 28dpc differentially expressed myelin proteins from the Siems et al. dataset are shown (n = 244 proteins). A Friedman-test was conducted, with a p-value ≤ 0.05. Post-hoc Dunn’s multiple comparison tests were conducted with * = p ≤ 0.05 for comparisons 7d crush versus intact and 28d crush versus intact and # = p ≤ 0.05 for the comparison 28d crush versus 7d crush.

### Proteomic analysis and quantification of mouse peripheral sciatic nerve myelin proteins

So as to determine the relative contribution of individual myelin proteins in young and old sciatic nerves, we calculated the relative abundance of each detected myelin protein as percentage of the total proteome, based on the mean iBAQ-values (i.e. the protein intensity within the corresponding sample) of the respective protein within each experimental group. Consistent with previous reports that provided accurate quantifications of the whole myelin proteome, the well-known myelin proteins MPZ, MBP and Periaxin constitute approximately 60-75% of the whole myelin proteome; ∼63% in young myelin, ∼74% in old myelin. (**Figure 3A, B**). Whilst the absolute protein abundances of MPZ and MBP did not change in old myelin, their relative abundance within the whole myelin proteome increased slightly (MPZ from 40 to 49%, MBP from 17 to 20%), which may be related to a general tendency of all detected myelin proteins to have lower protein abundancies in young as compared to old nerves (**Figure 3C** and **4**). However, the majority of myelin proteins were detected in both old and young nerves (∼90%) and the overall protein abundancies in young and old myelin fractions are closely correlated, suggesting no major changes in isolated myelin fractions during PNS aging (**Figure 3C**).

**Figure 3.**
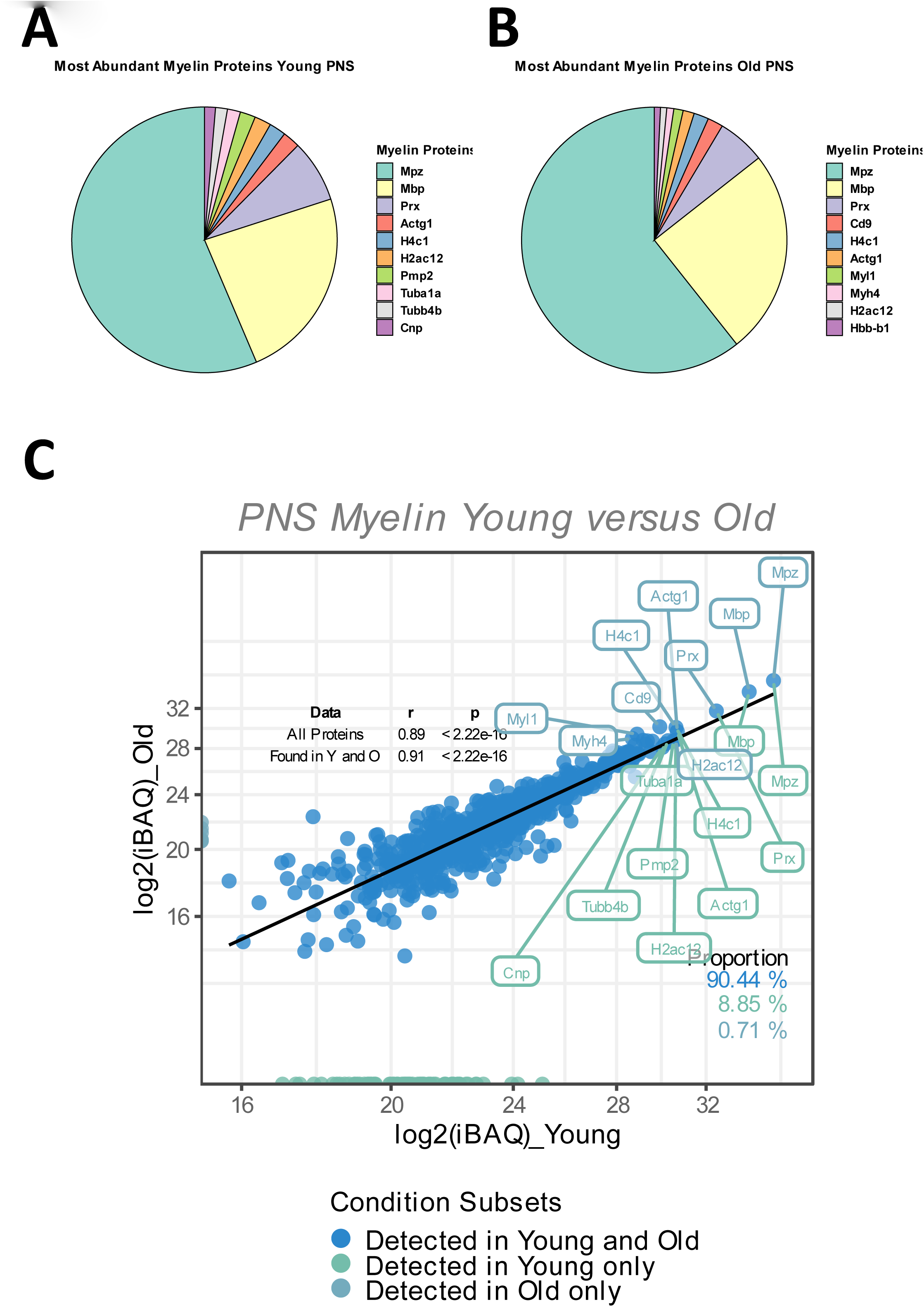
**A, B** Pie chart of the relative abundance/iBAQ-values of myelin proteins detected in extracted sciatic nerve myelin from young (**A**) and old (**B**) mice (n = 3 nerve pools of 7 to 10 nerves) as detected via mass spectrometry. **C** Correlation plot of the mean log_2_(iBAQ-values) of all myelin proteins detected in young and old sciatic nerves. All proteins detected in young and old nerves are coloured in blue. Those detected only in young nerves are labelled in green, with all proteins detected only in old nerves labelled in turquoise. The percentage of proteins for each subfraction is shown in the lower right corner. The correlation coefficient (Spearman’s ρ) and significance of all proteins, as well as proteins that are found in either young or old nerves, are shown in the upper right corner. The 10 proteins with the highest mean iBAQ-values in each fraction are labelled with their corresponding gene names (the top 10 young proteins in green, the top 10 old proteins in turquoise).

**Figure 4.**
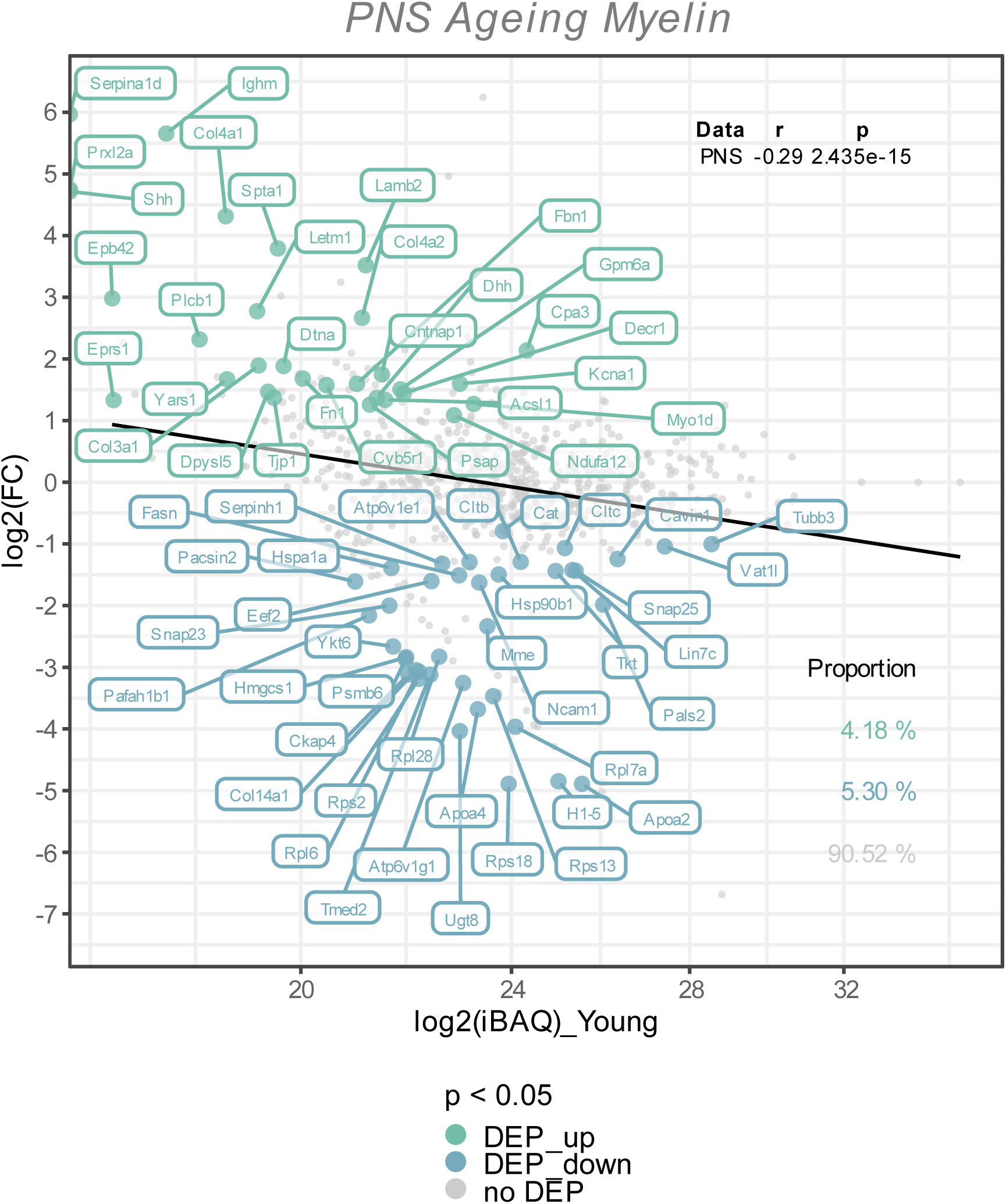
The Log_2_ FC from the old versus young myelin proteome comparison is plotted against the Log_2_ iBAQ-values, to visualize the differentially expressed proteins in relation to their abundance rank, i.e., the iBAQ-value in the young myelin proteome dataset. All differentially expressed proteins (FDR tool p-value < 0.05) are coloured and labelled; in old nerves upregulated myelin proteins in green, downregulated proteins in turquoise. The proportion of these subsets are shown in the lower right corner, with correlation statistics in the upper right corner.

### Proteomic changes in PNS myelin from old mice

To visualize potential proteome changes in old PNS myelin fractions in relation to their abundance in young PNS myelin proteomes, we plotted the fold changes of all proteins from old to young myelin fractions and the mean iBAQ-values of the isolated young myelin (**Figure 4**). As indicated above, the vast majority of proteins did not change significantly, implying that myelin protein levels are generally stable in isolated myelin from old nerves. However, correlation analysis of the entire proteome, including all non-significant proteins, revealed a negative association between fold changes and protein abundance in young nerves, pointing toward a general tendency for highly abundant myelin proteins to be downregulated during aging – although not significant at the individual protein level (**Figure 4**). This observation is mirrored in the differentially expressed proteins (DEPs): Downregulated proteins tended to be more abundant than upregulated proteins (**Figure 4**). Interesting and unexpectedly identified DEPs amongst all downregulated DEPs included e.g., several ribosomal proteins (Rps18, Rps13, Rps2, Rpl7a) and Ugt8, one of the major myelin lipid synthesizing enzymes (43). Several extracellular proteins were identified in the group of upregulated proteins, such as collagens (Col4a1, Col4a2, Col3a1, Fbn1) and other proteins relevant for extracellular matrix (ECM) interactions (e.g. Lamb2, Cntnap1). For an unbiased analysis of those biological pathways the up-and downregulated proteins are associated with, we performed an enrichment analysis for Reactome pathways (**Figure 5A**). The majority of pathways enriched in the list of downregulated proteins were related to ribosomal translation events. Furthermore, myelin proteins upregulated with age are mainly associated with ECM interactions and ECM organization, particularly collagen formation (**Figure 5A**). Therefore, all proteins were mapped to their gene ontology term (GO-term) cellular component annotations. Validated extracellular matrix components were both down-and upregulated – slightly more proteins were upregulated than downregulated – with several DEPs among the upregulated proteins, in particular several collagen proteins. Of these, Col4a1, a known peripheral nerve ECM component relevant to myelin organization (44), was the most upregulated protein (Log_2_FC = 4.32, **Figure 5B**).

**Figure 5.**
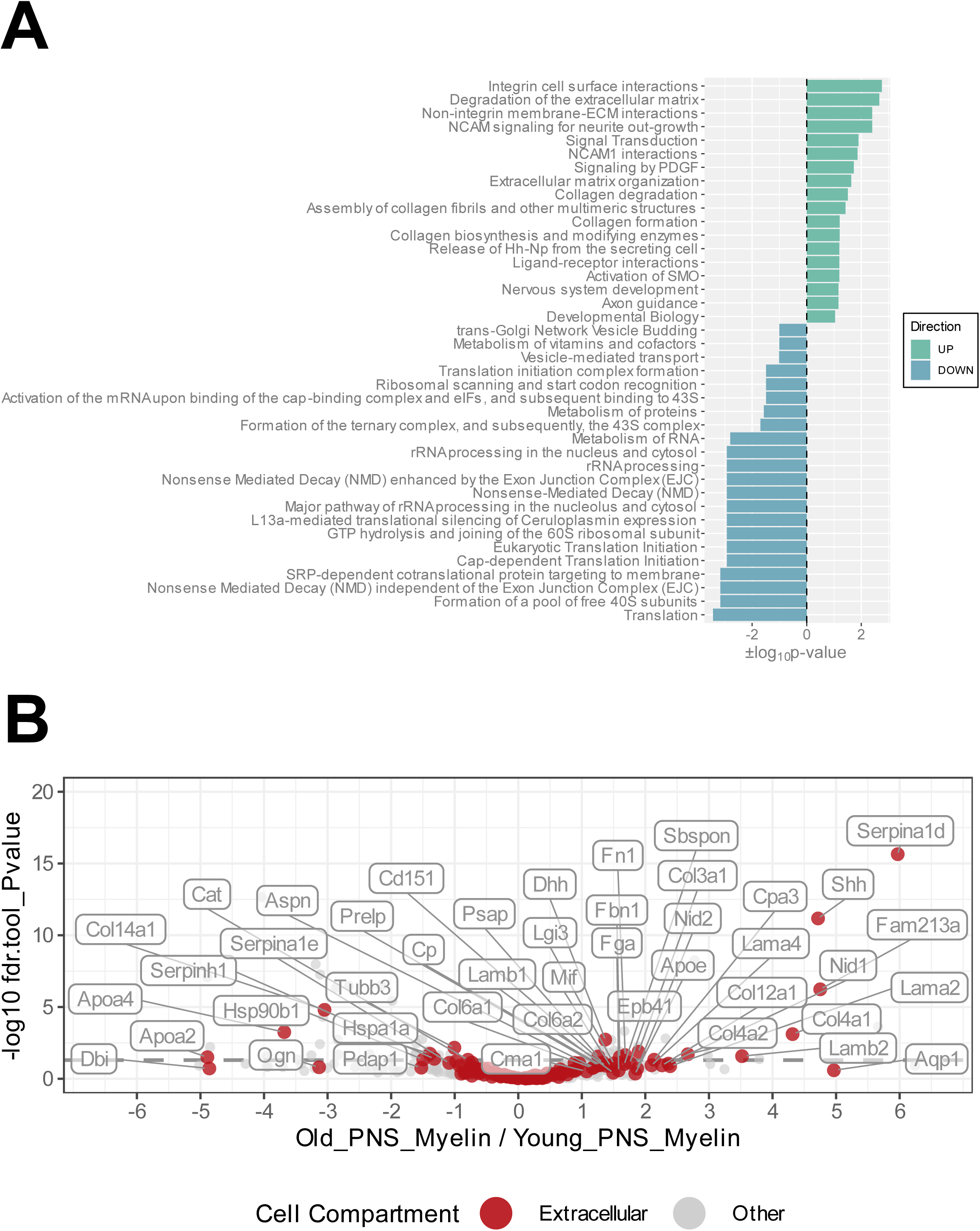
**A** Functional Reactome pathway analysis of DEPs was conducted using the DAVID platform. Pathways with a Fisher Exact p-value ≤ 0.1 were considered to be significantly enriched. Pathways found to be enriched in the list of the upregulated myelin proteins are labelled in green and ranked according to their positive log_10_p-value; pathways found to be enriched in the list of the downregulated myelin proteins are labelled in turquoise and ranked according to their negative log_10_p-value. **B** Volcano plot depicting the Log_2_FC and p-values for all detected and quantified myelin proteins. All gene ontology cellular component annotations were retrieved for all myelin proteins; those annotated to extracellular compartments are labelled in red. In addition, extracellular proteins that were differentially expressed (p-value < 0.05) are labelled with their respective gene names.

### Immunohistochemical validation of Col4a1 upregulation in the old PNS

Proteomic analysis predicted an upregulation of Col4a1 in myelin fractions isolated from old mouse sciatic nerves. This finding was validated by conducting immunofluorescence stainings on young and old transverse sciatic nerve sections (**Figure 6**). Col4a1 fluorescence intensity increased by ∼48% in total in old nerves, representing a very large effect of aging on Col4a1 protein levels in old sciatic nerves (Cohen’s d = 4.8) (**Figure 6A** and **C**). A closer examination of Col4a1 staining further revealed that endoneurial collagen showed considerable overlap with myelin sheaths (MBP immunostaining) in old nerves, in contrast to young nerves (**Figure 6B**). Quantification of Col4a1 myelin coverage (i.e., the percentage of Col4a1^+^ area within the total MBP^+^ area colocalized with MBP signal) also revealed increased overlap of Col4a1 fluorescent signal with MBP signal, potentially indicating greater proximity of basal lamina type IV collagen to old myelin (**Figure 6D)**. We also found that the fluorescence intensity of Col4a1 in the Col4a1/MBP overlapping endoneurial area increased in old nerves, suggesting not only a redistribution but also increased collagen levels adjacent to old PNS myelin compartments (**Figure 6E**).

**Figure 6.**
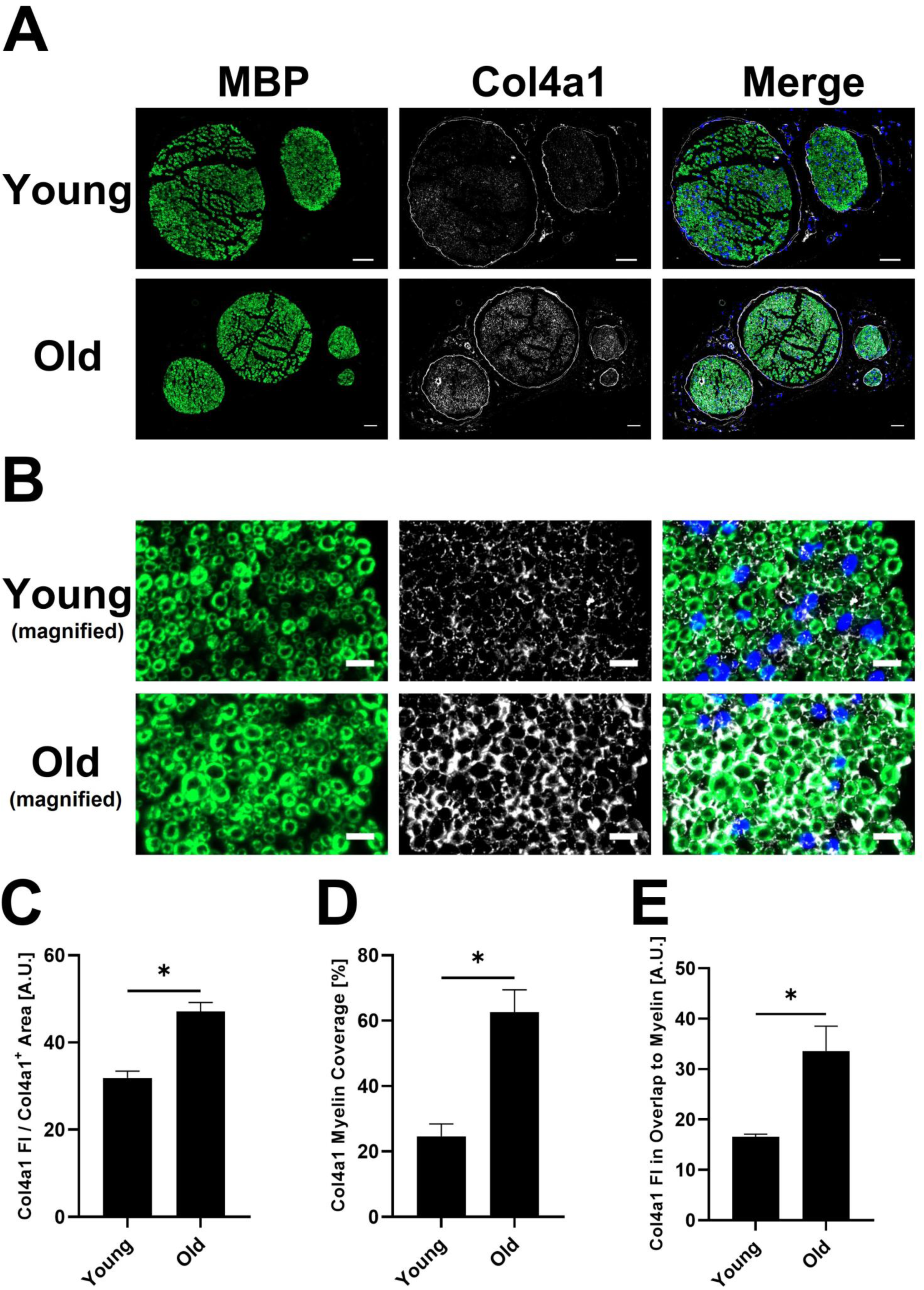
**A** Representative transverse whole sciatic nerve sections of young and old mice stained for the myelin basic protein (MBP) (A488 (green), left image) and the in the old myelin proteome strongest upregulated collagen subtype Col4a1 (A647 (white), middle image), as well as merged images of both channels in addition to the DAPI channel (right image), imaged on an Axio Scan.Z1 device with a 20x objective. Scale bars = 50µm. **B** Magnified image sections from **A** show overlap between myelin and collagen signals. MBP staining on the left, Col4a1 in the middle, merged channels with DAPI channel on the right. Scale bars = 10µm. **C** Quantification of the Col4a1 fluorescence intensity in young and old sciatic nerves normalized to the total Col4a1^+^ area in the respective image, divided by 10^5^ (n = 3 nerves per group, mean ± SEM, * = p<0.05, unpaired Student’s t-test.) **D** Quantification of the Col4a1^+^ area within the MBP^+^ area in %, i.e., percentage of the total Col4a1^+^ area that overlaps with MBP fluorescent signal in young and old sciatic nerves (n = 3 nerves per group, mean ± SEM, * = p<0.05, unpaired Student’s t-test.) **E** Quantification of the Col4a1 fluorescence intensity normalized to the Col4a1 myelin coverage and MBP^+^ area in young and old sciatic nerves, divided by 10^3^ (n = 3 nerves per group, mean ± SEM, * = p<0.05, unpaired Student’s t-test.)

## IV. Discussion

Aging brings multiple changes in peripheral nerve function, histopathology and molecular pathways (1, 2, 3, 17, 45). Dysfunctional myelin clearance and morphological alterations of myelin itself have been described in aged nerves (2, 4), which may contribute to aging nerve pathology. However, changes in peripheral nerve myelin itself during aging have yet to be investigated. To this end, we isolated PNS myelin from young and old mice to describe age-related changes in peripheral nerve myelin, extending previous studies describing the PNS myelin proteome in health and disease, such as neuropathies (12, 13).

As previously reported (13), we confirm that the three major myelin proteins MPZ, MBP and PRX constitute the majority of peripheral nerve myelin in both young and old nerves (63% in young, 74% in old nerves, **Figure 3**). Surprisingly, we detected only subtle changes in the total myelin proteome during PNS aging at the level of individual proteins (**Figure 4**). This is in contrast to the strong myelin remodeling observed in the Prx^-/-^ neuropathy mouse model, where approximately 30% of the proteome was seen to be differentially altered (q < 0.05)(13). Nevertheless, we found an overall negative correlation for fold changes in old versus young myelin and relative protein abundance in young myelin, suggesting a possible decrease in highly abundant myelin proteins other than MPZ, MBP and PRX; leading to a relative increase of these proteins in old myelin, despite their overall protein abundance remaining unchanged (**Figure 3C** and **4**). This observation echoes that observed in the Prx^-/-^ mouse model: PRX knockout resulted in an increased relative abundance of MBP and MPZ “to fill the gap” left by PRX loss (13). In addition, we found that the majority of previously accurately quantified PNS myelin proteins (13) were also detected in our proteomics dataset (**Figure 1A**) and that the protein quantities correlated significantly and positively with the relative quantities reported by Siems et al. (**Figure 1B**), indicating a qualitatively good myelin proteome as reported in this study. Furthermore, we wanted to test whether the determined myelin proteins are also downregulated during peripheral nerve degeneration, as myelin typically becomes degraded and cleared after a peripheral nerve injury (2, 37, 42). Myelin clearance is initiated within hours post-injury (37, 46) with myelin protein removal largely completed within 7 to 10dpc (42). Consistent with this was our observation that the majority of myelin proteins identified were significantly downregulated in the total nerve proteomes at 7dpc (**Figure 2**) – further evidence that the determined proteomes of isolated myelin fractions are indeed associated with myelin. Besides that, we detected that the myelin proteins that were downregulated at 7dpc, became upregulated again at 28dpc (**Figure 2**); which can be considered a classic sign of regeneration, as it is known that remyelination and nerve regeneration are already quite progressed 4 weeks post injury (17). Therefore, we contend that the peripheral myelin proteome reported in this study – but also the overlap with previously determined myelin proteome profiles (**Figure S1**)(13) – can on the one hand be used in unbiased proteomics studies to accurately quantify myelin clearance and degradation during e.g. peripheral nerve de-and regeneration, and on the other hand to detect differences in myelin removal in e.g. different pathological mouse models.

Although no major remodeling of the peripheral nerve myelin proteome was detected, we identified subtle changes in specific proteins to be differentially abundant in old versus young myelin fractions (**Figure 4**). Reactome pathway enrichment analysis revealed that significantly downregulated myelin proteins were mainly associated with translation mechanisms and ribosomes (**Figure 4**, **5A**). The identification of mainly ribosomal proteins associated with myelin was surprising as it is difficult to imagine any function for ribosomes as myelin constituents. However, almost all ribosomal proteins identified in our proteomics experiment were also identified in the previously reported PNS myelin proteome by Siems et al.(13) – suggesting that ribosomal proteins are indeed a true constituent of at least the myelin fraction that can be isolated from peripheral nerves. Of interest here is that it has previously been reported that Schwann cells can transfer ribosomes and mRNA to axons via narrow channels within myelin sheaths, the so-called Schmidt-Lantermann incisures (SLI), both under homeostatic conditions (47) and after injury (48). This Schwann cell/axonal transport of ribosomes is crucial to regeneration of injured axons; a previous study reported a reduced number of axonal ribosomes during aging, which may contribute to the reduced axonal regenerative capacity during aging (49). We speculate that vesicle-packed ribosomes within SLIs may be isolated together with the myelin due to entrapment within myelin particles, without actually being a component of myelin. The reduced quantity of ribosomal proteins identified in old myelin fractions may indeed be related to a reduced transfer of ribosomes from Schwann cells to axons in aged nerves, which may contribute to age-related axonopathy and reduced regenerative potential. Age-related reductions in ribosomes and translation and loss of proteostasis have been linked to aging in multiple organs (50) and are now considered a hallmark of aging (51). Yet, these speculative assumptions demand further study to measure translational activity and ribosome protein expression in Schwann cells during aging, as well as potentially altered Schwann cell to axon ribosome transfer.

Upregulated proteins were mainly enriched in pathways related to extracellular matrix organization, with several ECM-proteins, in particular collagens, induced in aged nerves (**Figure 4, 5**). Similar to the identified ribosomal proteins, collagens have been reported by Siems et al. to be part of isolated myelin fractions (13). Collagen proteins are integral components of the Schwann cell basal lamina (“Schwann tube”)(52) and thus mainly constitute the endoneurium in close proximity to and continuously surrounding myelin-wrapped axons (52, 53). In fact, the most abundant basal lamina collagens in peripheral nerves are heterotrimers of type IV collagens Col4a1 and Col4a2, produced by Schwann cells (53, 54). During development, type IV collagen is essential to formation of peripheral nerves and the induction of myelination (44, 55). However, these are known to increase during aging in humans (56), in neuropathies (57, 58, 59) and after PNS injury (60, 61), forming so called “collagen pockets” that can be wrapped by Schwann cell processes similar to axon myelination (62). In addition to their morphological functions, collagens can influence Schwann cell behaviors such as migration, differentiation and myelination (44, 53). Inhibition of endoneurial type IV collagen deposition has been linked to improved axon regeneration in the CNS (63), illuminating a potential regeneration-constraining effect of increased type IV collagen in the old PNS. Importantly, in a mouse model of neurofibromatosis type 2 (NF2), in which regeneration after a crush injury is strongly impaired, type IV collagen was found to be markedly increased post-injury compared to wildtype nerves (60). We found that not only was type IV collagen generally increased in old sciatic nerves, but also that these collagens were more likely to be adjacent to myelin (**Figure 6**). This may be indicative of thickening of the basal lamina of Schwann cells surrounding myelinated axons, leading to increased overlap with immunostained myelin/MBP, as observed in our fluorescence microscopy images. This thickening of the basal lamina with advanced age (64) and increased abundance of collagen pockets and their encirclement by Schwann cells in aged nerves (1), have been described previously and tally with our results. However, whether collagens can actually bind to myelin sheaths during aging is doubtful and merits further investigation, e.g. by immunogold labelling and electron microscopy. Nevertheless, we have shown increased endoneurial basal lamina-associated type IV collagen in aged nerves, which may have potentially negative effects on Schwann cell function and regenerative capacity.

In conclusion, we described for the first time the proteomic composition of peripheral nerve myelin in old mice. We showed that the 3 most abundant peripheral myelin proteins (MPZ, MBP and PRX) do not change with age in isolated myelin fractions, but high-abundance myelin proteins tended to decline, although we did not identify significant changes at the individual protein level for most of the proteins detected. Proteins that were found to change significantly in old myelin isolates were mainly ribosomes and extracellular proteins, such as collagens. These differences may reflect age-associated changes in the peripheral nerve rather than changes in the myelin composition itself. We have further shown that the myelin proteome determined in this study can be used to unbiasedly assess myelin degradation and replenishment during peripheral nerve de- and regeneration. All in all, we believe that this study can provide a resource for other researchers investigating aging myelin and the aging peripheral nervous system, as well as representing a starting point for further investigation of myelin remodeling during aging and after peripheral nerve injury.

## Acknowledgments

The authors would like to thank Rose-Marie Zimmer and Norman Rahnis for excellent technical assistance. Furthermore, the authors would like to thank Dr. Birgit Perner (Imaging Facility FLI Jena) for help in setting up an imaging protocol for the Axio Scan.Z1 device, Sabine Landmann-Weinsheimer for assistance with histological preparation of sciatic nerves, Madelaine Braune and Claudia Maisch for animal husbandry and Leonie Karoline Stabenow for critical reading and editing of the manuscript. Open access funding provided by Projekt DEAL.

